# Establishing a comprehensive web-based analysis platform for *Nicotiana benthamiana* genome and transcriptome

**DOI:** 10.1101/2023.09.03.556139

**Authors:** Ken-ichi Kurotani, Hideki Hirakawa, Kenta Shirasawa, Koya Tagiri, Moe Mori, Yasunori Ichihashi, Takamasa Suzuki, Yasuhiro Tanizawa, Yasukazu Nakamura, Sachiko Isobe, Michitaka Notaguchi

**Affiliations:** Bioscience and Biotechnology Center, Nagoya University, Nagoya, 464-8601, Japan; Department of Frontier Research and Development, Kazusa DNA Research Institute, Chiba 292-0818, Japan; Graduate school of Bioagricultural Science, Nagoya University, Nagoya, 464-8601, Japan; RIKEN BioResource Research Center, Tsukuba, Ibaraki 305-0074, Japan; College of Bioscience and Biotechnology, Chubu University, Matsumoto-cho, Kasugai 487-8501, Japan; Research Organization of Information and Systems, National Institute of Genetics, Yata, Mishima 411-8540, Japan; Department of Science, Kyoto University, Kyoto, 606-8502, Japan

## Abstract

*Nicotiana benthamiana* has long served as a crucial plant material extensively used in plant physiology research, particularly in the field of plant pathology, because of its high susceptibility to plant viruses. Additionally, it serves as a production platform to test vaccines and other valuable substances. Among its approximately 3.1 Gb genome, 57,583 genes have been annotated within a 61 Mb region. We created a comprehensive and easy-to-use platform to use transcriptomes for modern annotation. These tools allow to visualize gene expression profiles, draw molecular evolutionary phylogenetic trees of gene families, perform functional enrichment analyses, and facilitate output downloads. To demonstrate their utility, we analyzed the gene expression profiles of enzymes within the nicotine biosynthesis pathway, a secondary metabolic pathway characteristic of the *Nicotiana* genus. Using the developed tool, expression profiles of the nicotine biosynthesis pathway genes were generated. The expression patterns of eight gene groups in the pathway were strongly expressed in the roots and weakly expressed in leaves and flowers of *N. benthamiana*. The results were consistent with the established gene expression profiles in *Nicotiana tabacum* and provided insights into gene family composition and expression trends. The compilation of this database tool can facilitate genetic analysis of *N. benthamiana* in the future.

**Significance statement:** A tool was developed to visualize gene expression profiles, draw molecular evolutionary phylogenetic trees of gene families, perform functional enrichment analyses, and facilitate output downloads of *Nicotiana benthamiana*. The database developed using the tool can evolve into a comprehensive all-in-one analysis platform by continuously incorporating transcriptome data released to date, newly released RNA-seq data, and annotations in the future.

## Introduction

*Nicotiana benthamiana* is one of the oldest experimental models used in plant biology (Bombarely et al., 2012; Bally et al., 2018), and is valuable in plant pathology studies, notably, because of its susceptibility to a wide range of plant diseases, especially viral diseases. *N. benthamiana* has also recently received attention as a general platform for recombinant protein production. For example, the production of a COVID-19 vaccine has been achieved by inoculating and producing non-infectious virus-like particles (van Herpen et al., 2010; Bally et al., 2018; Mamedov et al., 2021). Interfamily grafting can also be realized in *N. benthamiana* (Notaguchi et al., 2020; Kurotani and Notaguchi, 2021). Although grafting is an old agricultural technique, its applicability has been limited, and it is considered to be established only among very closely related species in the same species or family. As *N. benthamiana* can achieve cell-cell adhesion at the graft junction with plants of different families, it has attracted attention as a target for studying self-recognition and wound repair mechanisms among plants.

*N. benthamiana* is a member of the family Solanaceae, which includes many agriculturally important crop species, such as potatoes, tomatoes, eggplants, petunia, and tobacco. *N. benthamiana* is native to Australia and almost 75 *Nicotiana* species are distributed in America and Australia. *N. benthamiana* in the *Suaveolentes* section forms n=19 allopolyploid, demonstrating that the paternal progeny belongs to the *Sylvestres* section (Knapp et al., 2004; Bally et al., 2018), whereas the maternal lineage may be the ancestor of the *Noctiflorae* section; however, the details are not clear. *Nicotiana tabacum*, the most widely cultivated and common tobacco species, is found in the *Nicotiana* section and descends from the ancestral species of the *Sylvestres* and *Tomentosae* sections. *De novo* whole-genome assembly was performed in *N. benthamiana* using HiFi reads, creating 1,668 contigs, 3.1 Gb in length. The 21 longest scaffolds considered pseudomolecules containing 2.8 Gb of sequence. In total, 57,583 high-confidence gene sequences were predicted within a 61 Mb region (Kurotani et al., 2023). Another study achieved construction of 19 pseudomolecules (Ranawaka et al., 2023). These two results are complementary to each other.

In this study, we aimed to create a usable database of the gene expression profiles of *N. benthamiana* based on stored genomic information. In addition to the previously conducted time-series transcriptomes of grafted plants, expression maps sampled by plant sites and site-specific transcriptomes of wound-treated or grafted plants were obtained to construct the database. By providing graphical expression profiles and a comprehensive set of analytical tools, including a sequence homology search, phylogenetic analysis, and heat mapping, it is expected that *N. benthamiana* will be more easily used as a model for plant science.

## Results and Discussion

### Database construction

To construct the *N. benthamiana* expression database, in addition to our previously published 11-point time-series transcriptome data of grafted plants, we performed transcriptome analysis of 28 new plant sites, yielding 117 RNA-seq datasets (Figure 1a, Supplementary Figure 1) of the stem apex, cotyledons, hypocotyls, and roots at the seedling stage; young and fully expanded leaf apical regions; floral organs; stems; and roots at the mature plant stage. These parts selected as the key tissues and organs in the *N. benthamiana* life cycle. Site-specific sectioned tissue for stems when grafted or wound-treated to the stem, and RNA-seq data over time for previously examined grafted plants were also included (Notaguchi et al., 2020). All RNA-seq data were processed using the same pipeline to obtain transcripts per million (TPM) and raw count matrices. Principal component analysis (PCA) was performed for each of the three groups: intact plant samples at all growth stages, wound-treated and grafted 7-day stems (Figure 1b,c). The transcriptomes of each of the three replicates were plotted at approximate positions, suggesting that the analyses were performed correctly. The leaves, roots, stems, petioles, and floral organs each formed four populations, reflecting the direction of differentiation (Figure 1b). The shoot apical meristems of seedlings were possibly not strictly separated from the hypocotyl. Stem samples on Day 7 of wound treatment and grafting were mostly similar and were plotted in similar positions for each stem site (Figure 1c). The wound and grafting treatments may have been nearly identical in terms of the physiological responses. PCA result of the time-series transcriptome of the grafting sites showed a behavior consistent with the previous mapping to the Niben1.01 genomic reference; however, the movement of the plots along the time-series depicted a more continuous linearity (Supplementary Figure 2, Notaguchi et al., 2020). This may have been due to the annotation accuracy.

**Figure 1.**
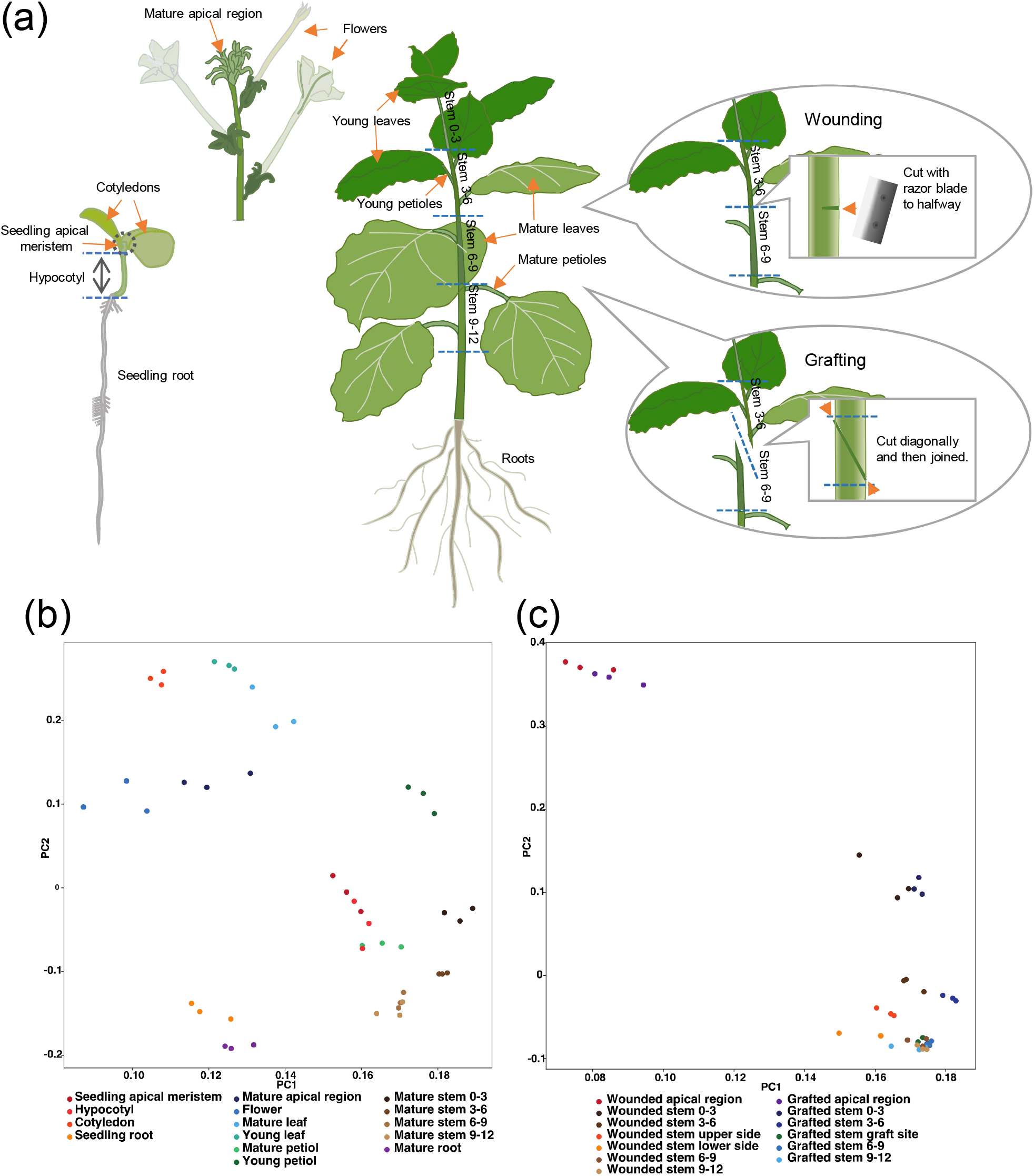
Transcriptome analysis by RNA-seq. (a) Plants and their parts used for RNA-seq analysis shown in the schematic illustration. (b,c) Principal component analysis for transcriptome. (b) Mature plants (7 weeks after germination) and seedling plants (7 days after germination). (c) Wounding or grafting (7 days after treatment).

The database provides tools for visualizing gene expression profiles, sequence homology searches, molecular phylogenetic tree generation from the obtained sequences, heat mapping and hierarchical clustering of expression profiles, and functional enrichment analysis (Figure 2). Each analysis tool allows the annotation list to be used as a hub for link analyses. Gene annotations can be searched using Nbe.v1 ID (Kurotani et al., 2023), Niben1.01 ID (Bombarely et al., 2012), and TAIR AT ID (Huala et al., 2001). A free-word search using gene description in the Arabidopsis annotation is also supported. By directly specifying a list of annotations or annotation IDs, the expression profile of each gene can be displayed as a color heat-pictograph (Figure 3). Simultaneously, users can display those profiles as bar graphs. The heat-pictograph can be displayed as either the absolute value of expression (TPM) or a normalized z-score, with the mean set to 0 and the standard deviation set to 1. The time-series transcriptomes of grafted stems are only available as a graph. Integrative Genomics Viewer (IGV, Robinson et al., 2011) was embedded to display the map status of sequence reads for all transcriptome data.

**Figure 2.**
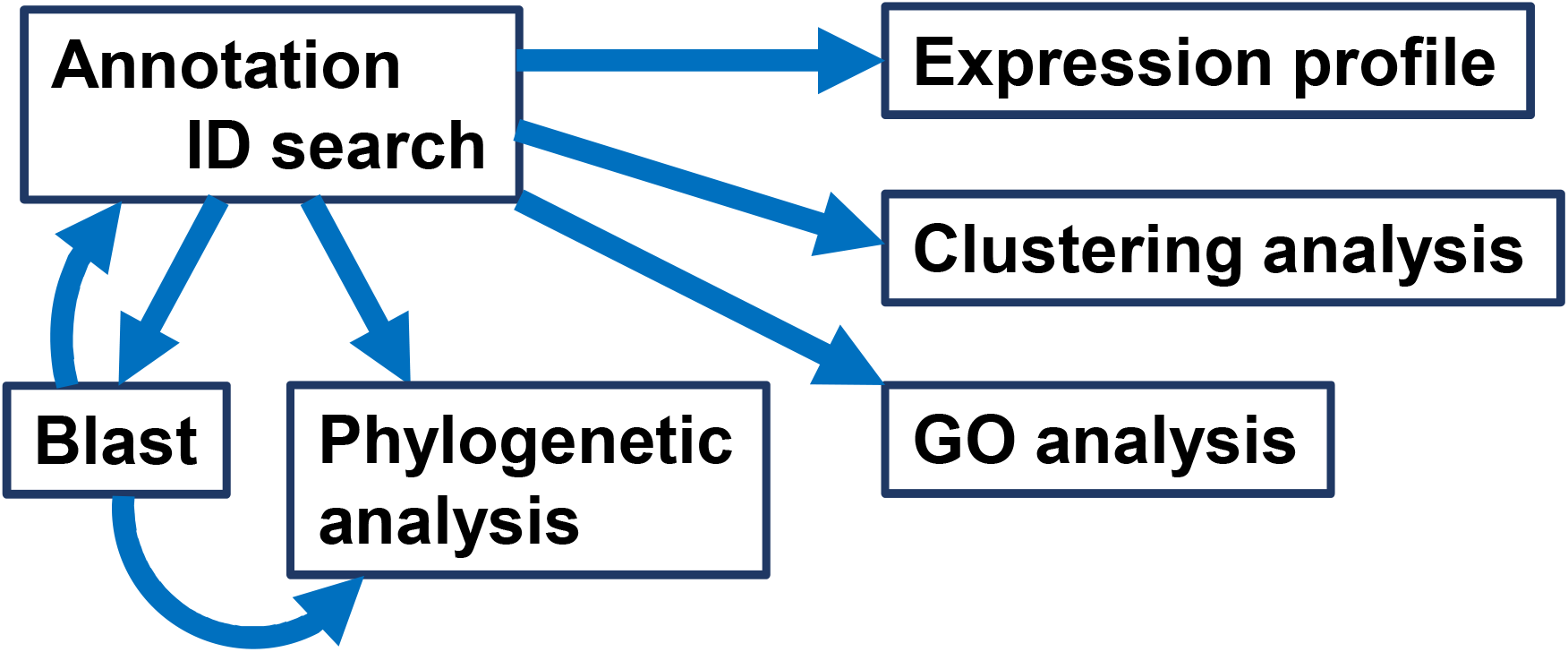
Database construction diagram.

**Figure 3.**
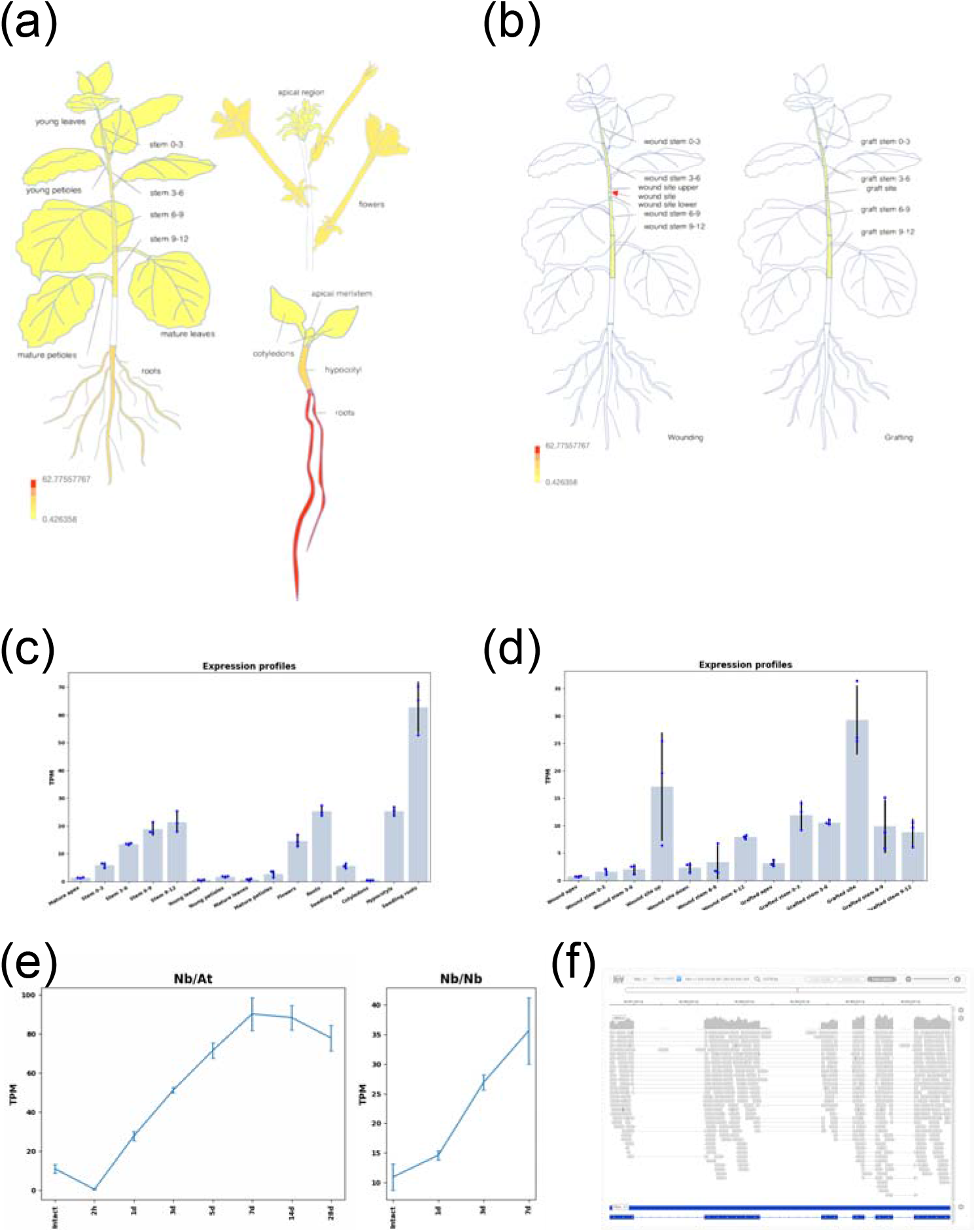
Construction of graphical expression browser. Examples of expression levels of *Nbe*.*v1*.*s00150g18250* gene. (a, b) Heat pictograph of gene expression levels (transcripts per million, TPM) by plant site in *N. benthamiana*. (a) Plants at 7 weeks after germination and seedling stages were separated by site. (b) Plants at 6 weeks after germination were wound-treated (left) or grafted, and the expression was examined in the stems 7 days after treatment. (c, d) Expression levels for three replicates are shown in the bar graph with error bars. (c) and (d) are bar graphs of (a) and (b), respectively. (e) Time-series transcriptome of grafted plants is displayed as a line graph. (f) The mapping status of the sequence reads from the RNA-seq analysis used to calculate the expression level is displayed in the Genome Browser. Error bars indicate standard deviation.

### Construction of gene search and sequence comparison tools

Blast+ of the National Center for Biotechnology Information (Camacho, 2009) was incorporated as a homology search tool for gene sequences. The coding sequence (CDS) sequence is obtained from the annotation of the gene to be examined, and the search can be directly applied to the BLAST program. The search targets are Nbe.v1 and Niben1.01 for *N. benthamiana*, and Araport11 of TAIR for *Arabidopsis thaliana*. Entire genome and transcript databases are available (https://nbenthamiana.jp). We implemented two search methods: blastn, which searched the DNA database using the DNA sequence as a query, and tblastn, which translated the query and DNA database into three frames of amino acids and searched only for transcripts. DNA sequences are entered directly as queries or FASTA format files are uploaded for analysis. The genes to be compared, obtained by BLAST, can be aligned with ClustalW using annotation IDs or their FASTA-formatted sequences, and a molecular phylogenetic tree can be drawn using the neighbor-joining method. The drawn images are provided in EPS format for easy processing and downloading (Supplementary Figures 3–5).

### Construction of gene expression analysis tools

A tool was prepared to create heat maps from the expression profiles of multiple genes, including gene families (Figure 4). This tool allows hierarchical clustering of genes from profiles. Clustering among samples was also made possible. The clustering algorithm can be selected from the Ward’s, single linkage, complete linkage, average linkage, weighted, centroid, and median methods. Distance can be selected from Euclidean, correlation, cosine, and city blocks. *N. benthamiana*, an allopolyploid species, has many genes in pairs that are located in the genome of two interbred species (Bally et al., 2018); therefore, similar genes often form large families. Because of this tool, these genes can be easily organized and evaluated, thereby facilitating genetic analyses, including comprehensive genome editing, using CRISPR.

**Figure 4.**
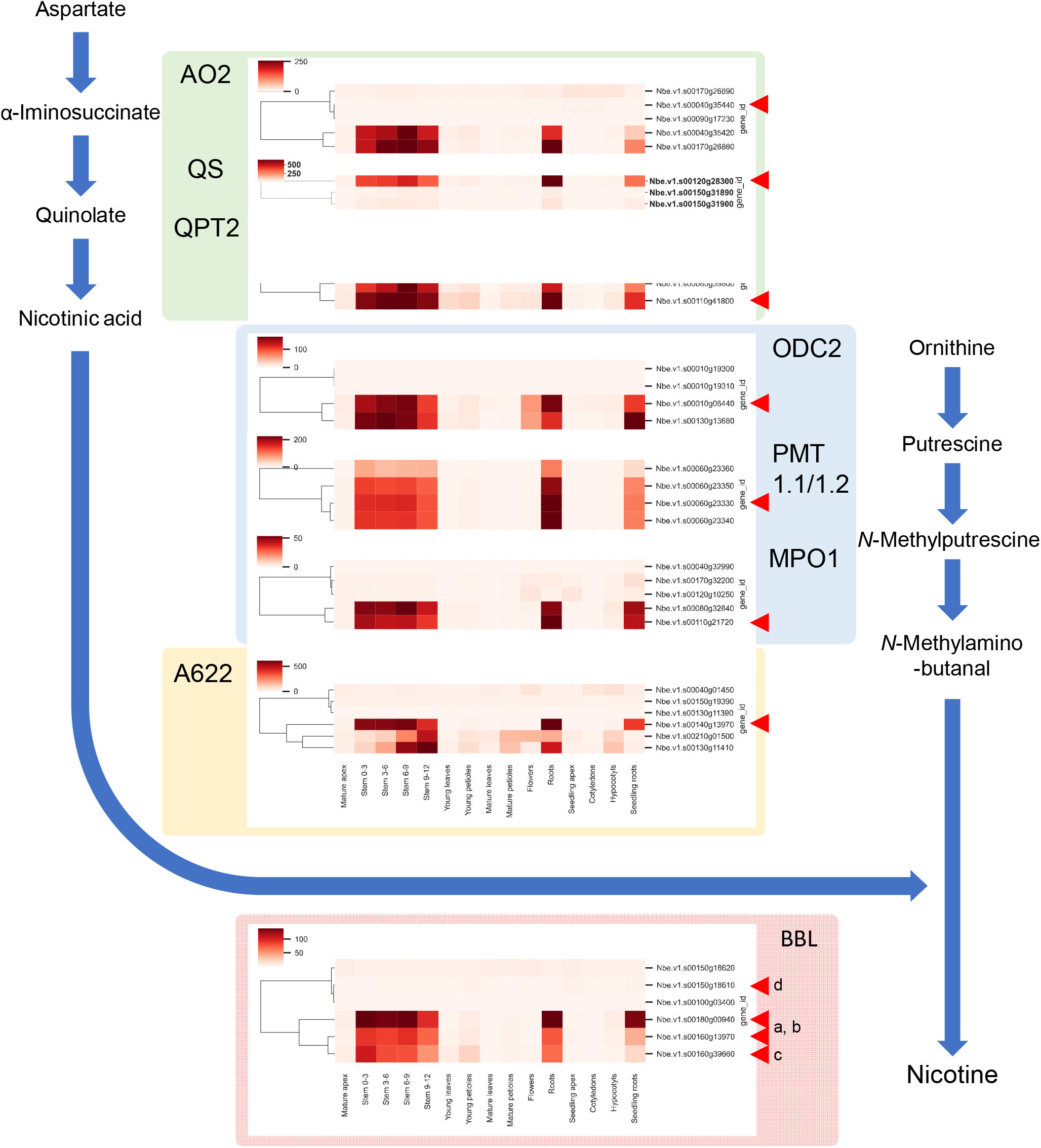
Expression analysis of a group of genes in the nicotine biosynthesis pathway. Profiles of *N. benthamiana* genes homologous to the eight genes involved in nicotine biosynthesis in *N. tabacum* were generated. Each gene is represented by a heat map, and a phylogenetic tree based on the expression profiles is placed to the left of the heat map. Genes that most closely approximated the *N. tabacum* genes are indicated by arrowheads.

### Case study: Expression profile analysis of nicotine biosynthesis pathway genes

To test the plausibility and utility of the developed database, we focused on the genes associated with nicotine biosynthesis in *N. benthamiana*. Nicotine is a major alkaloid and typological feature of plants of the genus *Nicotiana* and plays an important role in plant defense mechanisms (Hashimoto and Yamada, 1994; Steppuhn et al., 2004). Nicotine biosynthesis involves the pyrrolidine pathway metabolized from ornithine and the pyridine pathway metabolized from aspartate. Ornithine decarboxylase (ODC), putrescine *N*-methyltransferase (PMT), and *N*-methylptoresine oxidase (MPO) are involved in the formation of pyrrolidine rings (Hibi et al., 1994; Imanishi et al., 1998; Heim et al., 2007; Katoh et al., 2007). Aspartate oxidase (AO), quinolinic acid synthase (QS), quinolinic acid phosphoribosyltransferase (QPT), and other enzymes are involved in the early stages of nicotinic acid biosynthesis (Sinclair et al., 2000; Katoh et al., 2006) and nicotinic acid-derived precursors. The phosphatidylinositol phosphate (PIP) family oxidoreductase A622 and the berberine bridge enzyme-like protein (BBL) are suggested to be involved in the late bio-synthetic steps of pyridine alkaloids (Deboer et al., 2009; Kajikawa et al., 2009; Lewis et al., 2015; Vollheyde et al., 2023). Phylogenetic analysis suggests that the genes involved in the biosynthesis of pyridine and pyrrolidine rings evolved from the duplication of two major metabolic pathways that have long existed in all major plant lineages: the nicotinamide adenine nucleotide (NAD) coenzyme pathway and the polyamine metabolism pathway, respectively (Xu et al., 2017).

Transposons, which are characteristic of the Solanaceae family, have been suggested to cause the evolutionary doubling of these genes and their differentiation into pathways (NAD/pyridine ring pathways and polyamine / pyrrolidine ring pathways) with different regulation of expression and function (Xu et al., 2017). In the present study, we first examined the genetic components of the nicotine biosynthetic pathway genes in *N. tabacum* and their homologs in *N. benthamiana* (Supplementary Figures 3–5). Two *AO* genes were in the pyridine ring pathway, *NtAO1* and *NtAO2*, in *N. tabacum. NtAO2* was reported to be involved in nicotine biosynthesis, but three other *AO* genes were found in *N. tabacum*. However, two of these were extremely similar to *NtAO1* and may have been further diversified after doubling with *NtAO2*. The remaining gene was considered a partial double pseudogene without a complete gene. The homologs of *N. benthamiana* contained five genes. Two genes were close to *NtAO1* and the remaining three genes were close to *NtAO2*. This suggested that the separation of *NtAO1* and *NtAO2* occurred before *N. benthamiana* and *N. tabacum* were established. Interestingly, a pseudogene similar to *NtAO2* was also found in *N. benthamiana*, suggesting that this pseudogene might have been formed before the two species were established. The most similar genes to *NtAO1* and *NtAO2* were *Nbe*.*v1*.*s00040g35420* and *Nbe*.*v1*.*s00040g35440*, respectively. However, these two genes were located almost consecutively on the chromosome and were likely due to a relatively recent doubling; therefore, it may be more appropriate to consider *NtAO2* as *Nbe*.*v1*.*s00170g26890* (Supplementary Figure 3a). *QS* is believed to share a common gene that functions in the NAD and pyridine ring pathways. However, two copies of this gene are present in *N. tabacum*. The gene most closely related to *NtQSs* was *Nbe*.*v1*.*s00120g28300* (Supplementary Figure 3b). There were two *QPT* genes, *NtQPT1* and *NtQPT2*, in *N. tabacum*, and four genes were found in *N. benthamiana*; the genes most closely related to *NtQPT1* and *NtQPT2* were *Nbe*.*v1*.*s00080g39790* and *Nbe*.*v1*.*s00110g41800*, respectively (Supplementary Figure 3c). In the pyrrolidine ring pathway, on the other hand, only two *ODC* genes were found in *N. tabacum*, while four genes were found in *N. benthamiana*; the genes closely related to *NtODC1* and *NtODC2* were *Nbe*.*v1*.*s00010g19300* and *Nbe*.*v1. s00010g19310*, respectively. However, because these two genes are also located next to each other on the chromosome, we considered it reasonable to assign a homolog of *NtODC2* to *Nbe*.*v1*.*s00010g06440* (Supplementary Figure 4a). *PMT* were present in both species with five genes each; however, homologs of *NtPMT1* and *NtPMT2* could not be identified because the phylogenetic clade was split between *N. tabacum* and *N. benthamiana*. The five genes of *N. benthamiana* were arranged almost next to each other on the same chromosome, and their origin may not be consistent with the molecular evolution of *N. tabacum*. The closest homolog to *N. tabacum* was *Nbe*.*v1*.*s00060g23330* (Supplementary Figure 4b). *MPO* was found in *N. tabacum* with four genes, and in *N. benthamiana* with five genes. *NtMPO1* is involved in the nicotine biosynthesis pathway, but *PMO* genes are not involved in the polyamine pathway, and the homolog of *NtMPO1* was considered to be *Nbe*.*v1*.*s00110g21720* (Supplementary Figure 4c). Among the enzymes involved in the reaction that joins the two rings, A622 was found in two genes of *N. tabacum*. Similarly, five genes were found in *N. benthamiana*, but one homolog (*Nbe*.*v1*.*s00140g13970*), which was involved in the nicotine biosynthesis pathway, formed an independent clade (Supplementary Figure 5a). In *N. tabacum*, four *BBL* genes (*NtBBLa*-*NtBBLd*) may have been involved in this pathway. In *N. benthamiana*, there were six genes, and the homologs of *NtBBLa* and *NtBBLb* were *Nbe*.*v1*.*s00160g13970* or *Nbe*.*v1*.*s00180g00940*, respectively. Because they were in independent clades and their genetic distances were almost equal, distinguishing *BBLa* and *BBLb* was not possible. By contrast, the homologs of *NbBBLc* and *NbBBLd* were *Nbe*.*v1*.*s00160g39660* and *Nbe*.*v1*.*s00100g03400*, respectively (Supplementary Figure 5b).

The expression profiles of these nicotine biosynthesis pathway genes were generated using the tool developed in this study. The expression patterns of all eight gene groups were strongly expressed in the roots and weakly expressed in the leaves and flowers, as previously reported for *N. tabacum* (Xu et al., 2017). The hypothesis of the molecular evolution of nicotine biosynthetic pathways through gene doubling during the evolution to the *Nicotiana* genus in the Solanaceae family was also generally supported. The only difference was in the AOs, which are the primary enzymes in the pyridine ring pathway. In *N. tabacum*, NtAO2 metabolizes aspartate to α-iminosuccinate (Xu et al., 2017). However, in *N. benthamiana, AO* genes homologous to *NtAO1* were highly expressed in the roots, suggesting that the functional differentiation of these genes may have been acquired independently after establishing the two species.

### Conclusion and future prospects

The *N. benthamiana* database can perform a series of analyses, from annotation and sequence homology searches to web-based data visualization. This database is expected to evolve into a comprehensive all-in-one analysis platform by continuously incorporating transcriptome data released to date, newly released RNA-seq data, and annotations in the future. Currently, tools for differentially expressed gene analysis from expression data, expression network analysis, and pathway analysis have not yet been implemented. This database is expected to accelerate the molecular biological discovery of *N. benthamiana* and make *N. benthamiana* an easier-to-use plant research platform than ever before.

## Experimental Procedure

### Plant materials

*N. benthamiana* seeds were surface-sterilized with 5% (w/v) bleach for 5 min, washed three times with sterile water, incubated at 4°C for 3 d, and sown on half-strength Murashige and Skoog medium supplemented with 0.5% (m/v) sucrose and 1% agar. The pH was adjusted to pH 5.8 using 1 M KOH. *N. benthamiana* seedlings were grown at 27°C under continuous illumination intensity of 100 μmol m^-2^ s^-1^. On Day 7 after germination, the plants were replanted in a 1:1 mixture of soil medium and vermiculite. Stem grafting was performed as previously described (Notaguchi et al., 2020). Briefly, wedge grafting was performed on stems, and the grafted plants were initially grown in an incubator at 27°C under continuous light (ca. 30 μmol m^-2^ s^-1^) for a week and then transferred to a plant growth room at 22°C under continuous light conditions (ca. 80 μmol m^-2^ s^-1^). Stem wounding was performed under the same conditions as grafting without complete disconnection.

### Transcriptomic analysis

Plants growing to approximately 15 cm length 6–7 weeks after sowing were considered mature plants. Samples were taken from fully expanded mature leaves and their petioles, immature leaves and their petioles during elongation, stems cut from the stem apex every 3 cm, roots, flowering organs and immature inflorescence meristems. Three replicates of 10 plant samples per pool were used for the analysis. Cotyledons, shoot apical meristems, hypocotyls, and roots were sampled from the seedlings 7 d after sowing. Three replicate analyses were performed using one tissue pool from 20, 100, 100, and 25 individuals, respectively. Wound treatments and grafting were performed on the plants 4–5 weeks after sowing. For wound treatment, a razor blade was used to cut 6 cm from the stem apex to half the stem diameter, and sampling was performed after 7 d. The top and bottom of the wounds were located 1 cm above and below each cut, respectively, and remaining stems were removed every 3 cm. Similarly, during grafting, stems were cut diagonally 6 cm from the stem apex, joined using grafting clips, and sampled on Day 7. The grafted portion was sampled at a length of 1 cm without separating the upper and lower portions, and the remaining stems were cut into 3-cm pieces. Three replicates of 10 plants per pool were used for the analysis. Tissues were processed with zircon beads and lysate binding buffer containing sodium dodecyl sulfate instead of lithium dodecyl sulfate, as described in a previous report. RNA purification and cDNA library preparation followed the BrAD-seq method (Townsley et al., 2015). Further, 86-bp single-ended sequencing was performed on an Illumina NextSeq 500 platform (Illumina, San Diego, CA, USA). Data pre-processing was performed as follows. Data were trimmed for quality using Fastp v0.23.2 (Chen et al., 2018) with the settings qscore of 20 and length of 20. Trimmed reads were mapped onto the genome assembly using HISAT2 v2.1.0 (Kim et al., 2019). The generated sequence alignment and map (SAM) files were converted to binary alignment and map (BAM) format and merged using SAMtools v1.4.1 (Danecek, 2021). Gene expression levels (TPM) were estimated using StringTie v2.2.0 (Pertea et al., 2015).

### Browser tools

The backend of the expression database was implemented in Python using the Flask Web framework. Data were stored in SQlite3 database. The front end was developed using a bootstrap framework. The Integrative Genomics Viewer (IGV) was used to visualize the RNA-seq reads (Robinson et al., 2011). To normalize the expression levels, SciPy and scikit-learn libraries were used in Python (Pedregosa et al., 2012; Virtanen et al., 2020).

### Expression analysis and sequence comparison tools

The scipy.cluster.hierarchy.linkage, matplotlib, and seaborn.clustermap libraries were used for heat mapping and hierarchical clustering of expression levels (Hunter, 2007; Virtanen et al., 2020; Waskom, 2021). BLAST + was used to search for annotations or homologies in gene sequences (Camacho, 2009). Only DNA sequences were used for queries, and BLASTN and tBlastx were implemented. The search targets were Nbe.v1 and Niben1.01 for *N. benthamiana* and Araport11 for Arabidopsis, and both CDS or transcripts, and genomes were prepared. ClustalW2 (Thompson et al., 1994), which creates molecular phylogenetic trees from the annotation search results of multiple genes or FASTA format files containing multiple gene sequences, was incorporated. Google Library was used for gene ontology enrichment analysis from the annotation list (Klopfenstein, 2018).

## Data Statement

The genome assembly data, annotations and gene models, transcriptome data are available at the NbenBase (https://nbenthamiana.jp).

## Author Contributions

KK and MN conceived the research and designed the experiments. KK, HH, KS, KT, MM, YI, TS and YT performed the sample preparation and analyzed data with the support of YN, SI, and MN. KK and MN wrote the manuscript.

## Acknowledgements

We acknowledge technical assistance by Akiko Watanabe, Takaharu Kimura, Yoshie Kishida, Hisano Tsuruoka, Chiharu Minami, Akiko Komaki, Akiko Obara, Rie Aomiya, and Taeko Shibazaki of the Kazusa DNA Research Institute and Miki Matsumoto of Nagoya University.

## Supplementary figure legends

**Supplementary Figure 1. Plants and their parts used for RNA-seq analysis**

Bars:1 cm

**Supplementary Figure 2. PCA analysis for the time series transcriptome of interfamily grafting and homo grafting**.

*Nb*/*At* and *Nb*/*Nb* indicate interfamily grafting and homo grafting, respectively. RNA was extracted from grafted plants at 2 hours after grafting (HAG) and 1, 3, 5, 7, 10, 14 and 28 days after grafting (DAG).

**Supplementary Figure 3. Molecular phylogenetic trees of genes on the pyridine ring pathway**

(a) Aspartate oxidase (AO). (b) Quinolinic acid synthase (QS). (c) Quinolinic acid phosphoribosyltransferase (QPT).

**Supplementary Figure 4. Molecular phylogenetic trees of genes on the pyrrolidine ring pathway**

(a) Ornithine decarboxylase (ODC). (b) Putrescine *N*-methyltransferase (PMT). (c)

*N*-methylptoresine oxidase (MPO).

**Supplementary Figure 5. Molecular phylogenetic trees of genes on the late bio-synthetic steps of pyridine alkaloids**

(a) Phosphatidylinositol phosphate (PIP) family oxidoreductase A622. (b) Berberine bridge enzyme-like protein (BBL).

