## Supplementary Figures for "Establishing a comprehensive web-based analysis platform for *Nicotiana benthamiana* genome and transcriptome"

### Slide 1
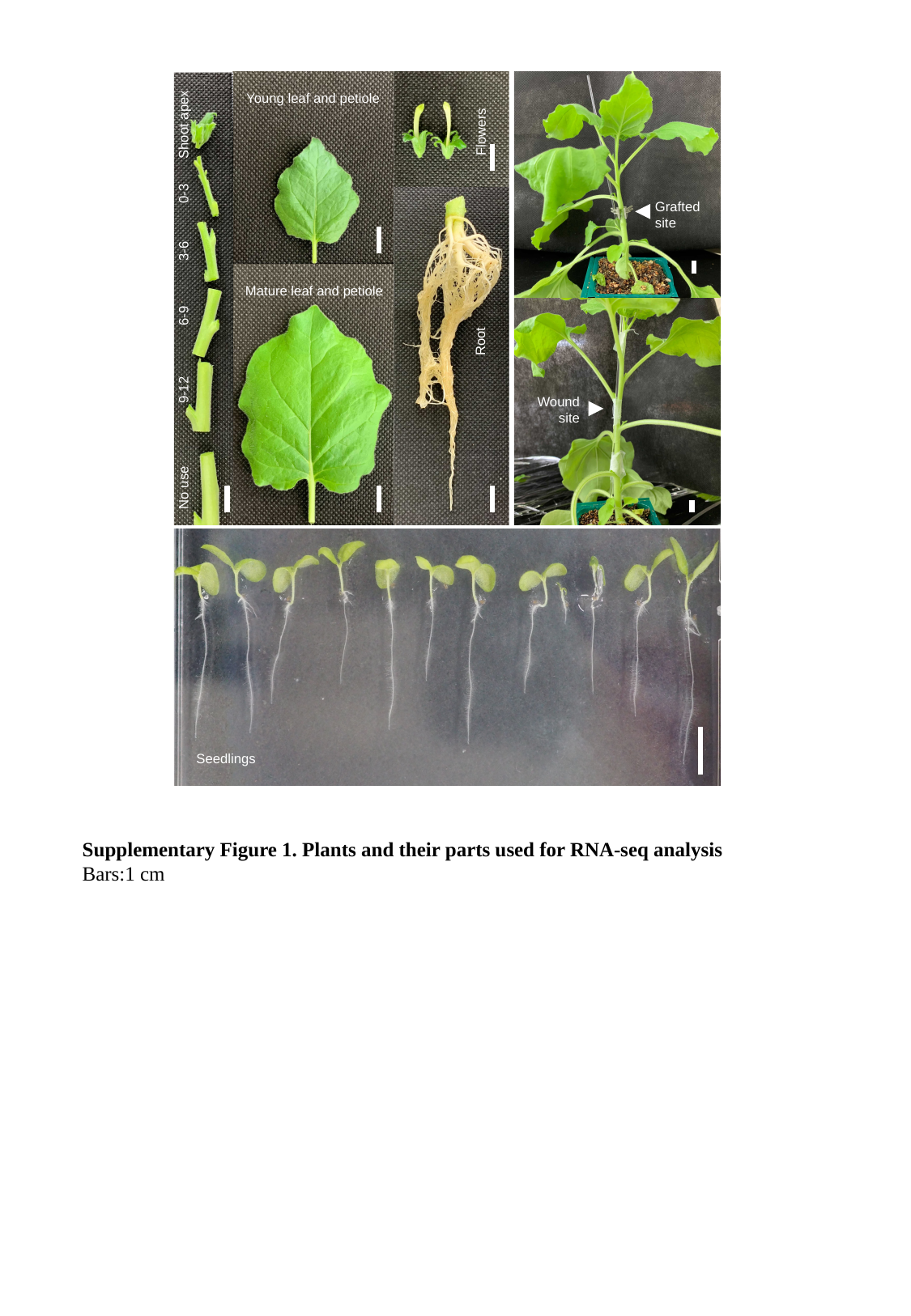

Young leaf and petiole
Shoot apex
Flowers
0-3
Grafted
site
3-6
Mature leaf and petiole
6-9
Root
9-12
Wound
site
No use
Seedlings
Supplementary Figure 1. Plants and their parts used for RNA-seq analysis
Bars:1 cm

### Slide 2
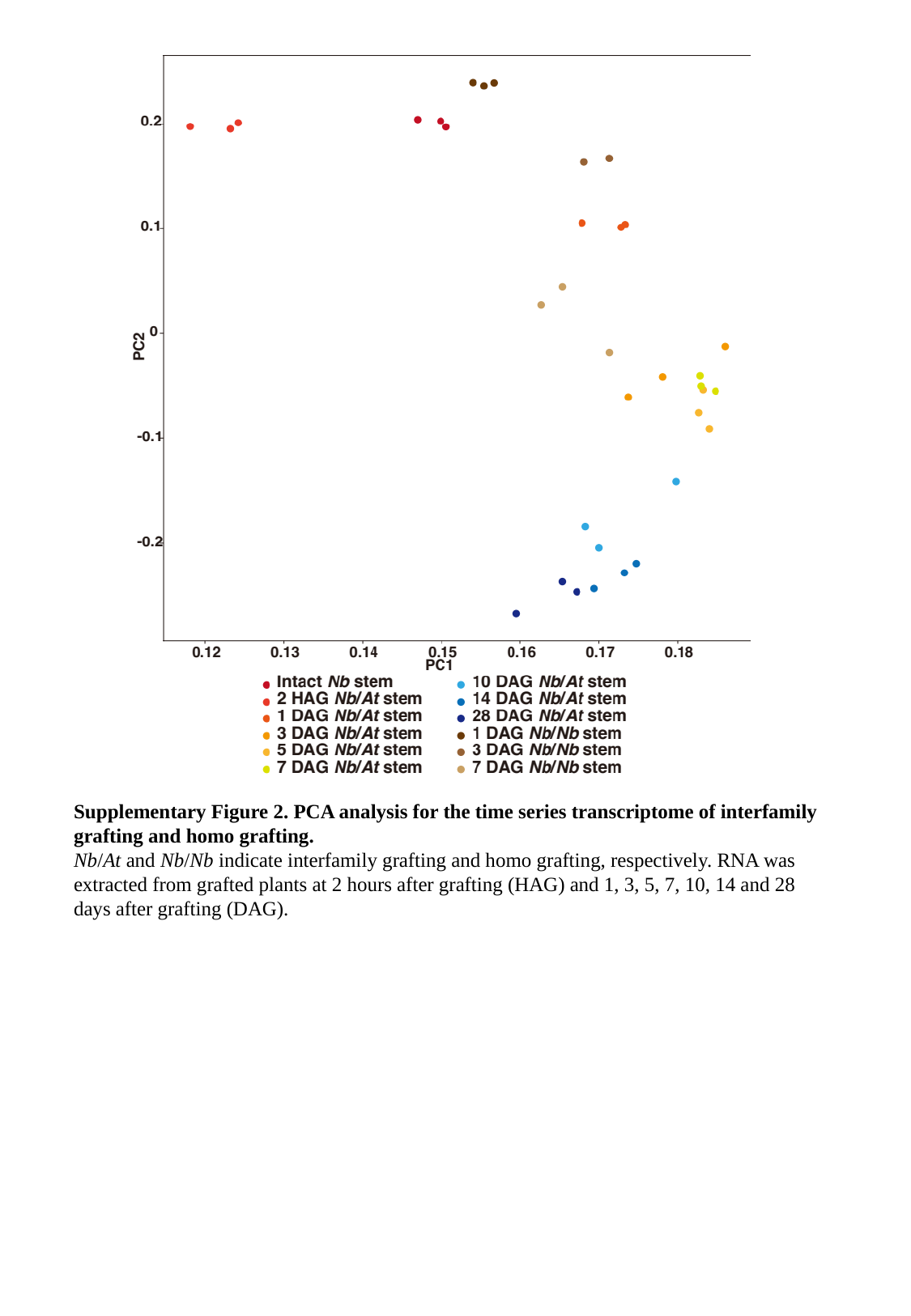

Supplementary Figure 2. PCA analysis for the time series transcriptome of interfamily grafting and homo grafting.
Nb/At and Nb/Nb indicate interfamily grafting and homo grafting, respectively. RNA was extracted from grafted plants at 2 hours after grafting (HAG) and 1, 3, 5, 7, 10, 14 and 28 days after grafting (DAG).

### Slide 3
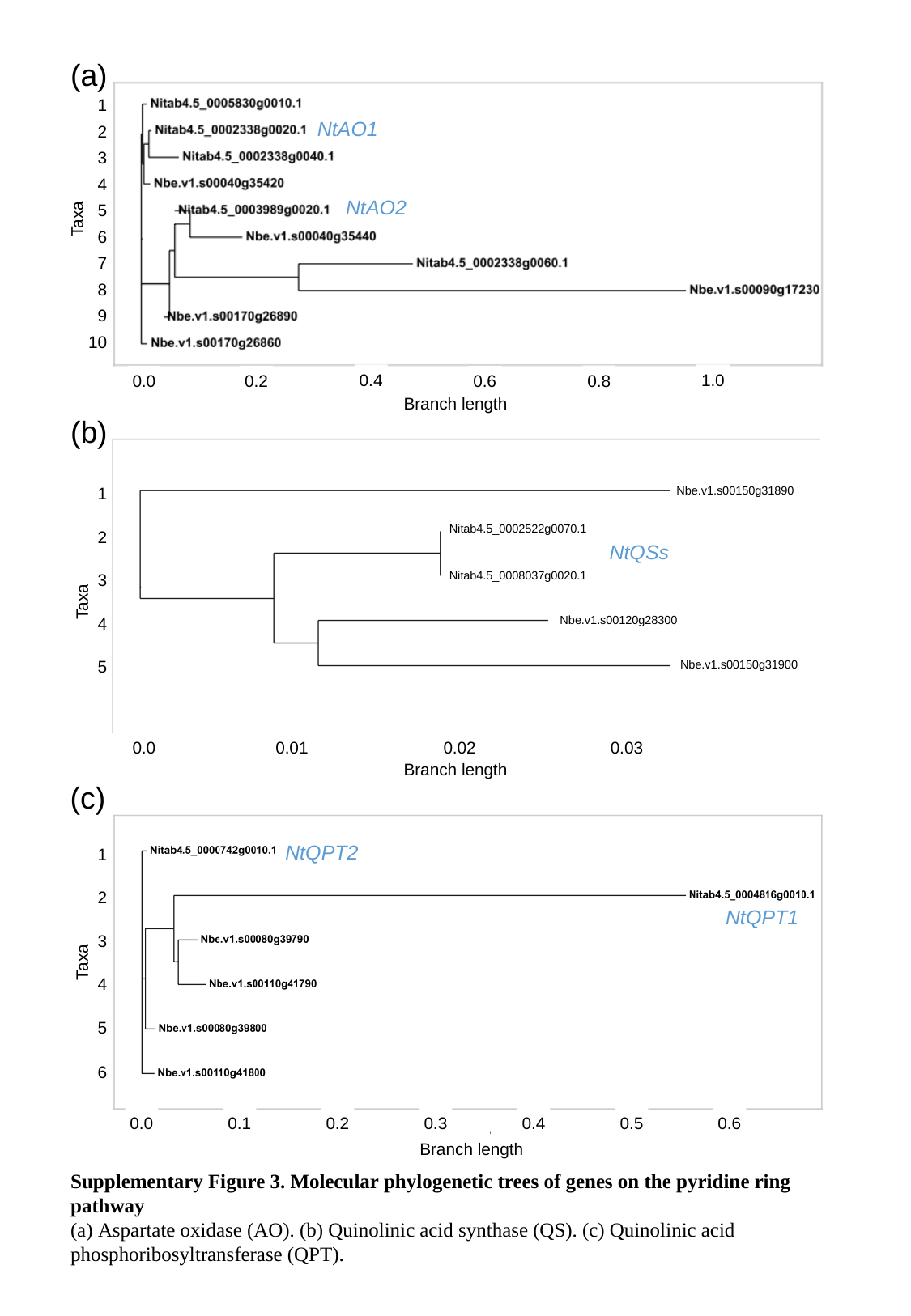

(a)
1
NtAO1
2
3
4
NtAO2
5
Taxa
6
7
8
9
10
0.4
1.0
0.8
0.2
0.6
0.0
Branch length
(b)
Nbe.v1.s00150g31890
1
Nitab4.5_0002522g0070.1
2
NtQSs
Nitab4.5_0008037g0020.1
3
Taxa
Nbe.v1.s00120g28300
4
5
Nbe.v1.s00150g31900
0.0
0.01
0.02
0.03
Branch length
(c)
NtQPT2
1
2
NtQPT1
3
Taxa
4
5
6
0.0
0.1
0.2
0.3
0.4
0.5
0.6
Branch length
Supplementary Figure 3. Molecular phylogenetic trees of genes on the pyridine ring pathway
(a) Aspartate oxidase (AO). (b) Quinolinic acid synthase (QS). (c) Quinolinic acid phosphoribosyltransferase (QPT).

### Slide 4
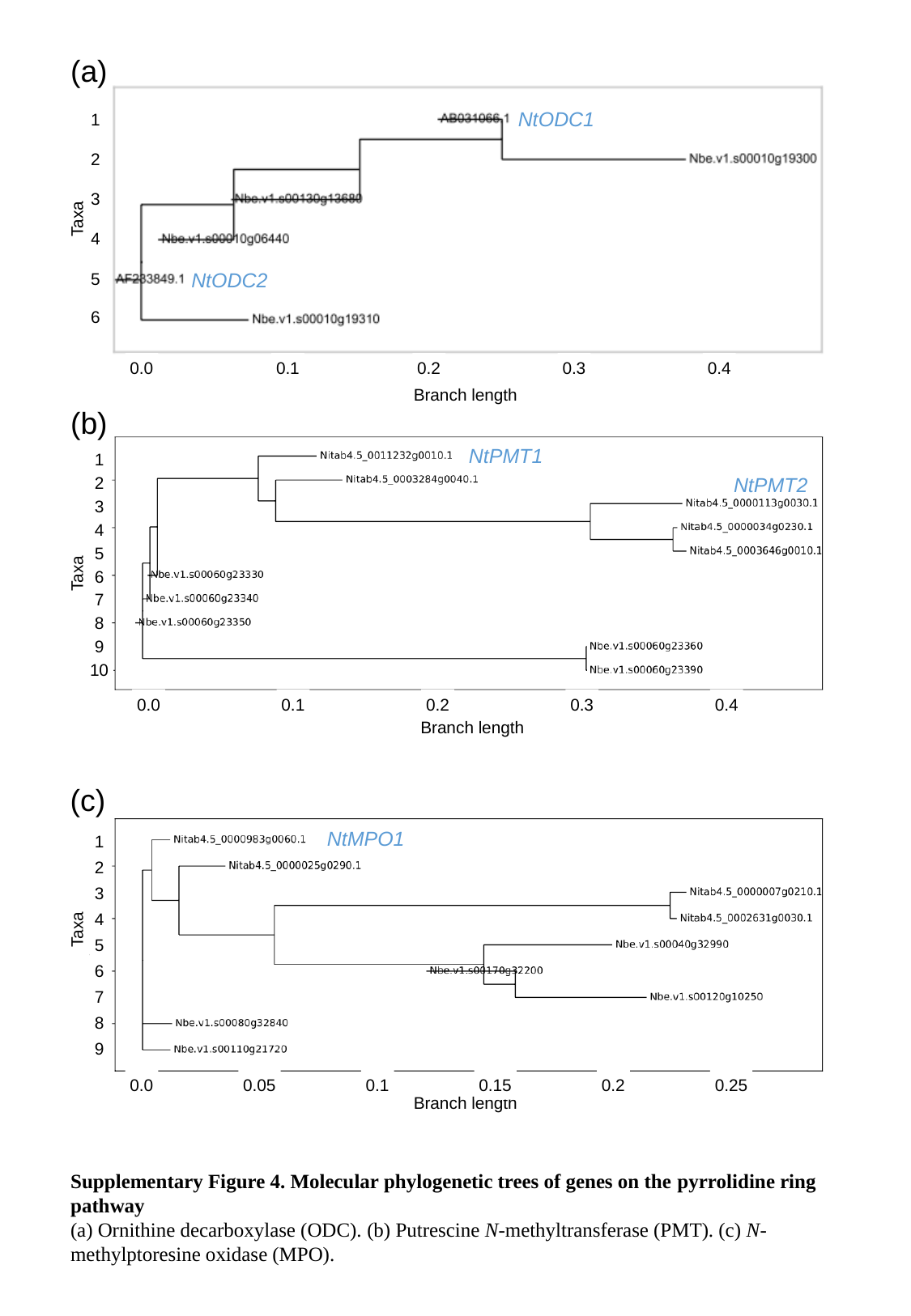

(a)
NtODC1
1
2
3
Taxa
4
NtODC2
5
6
0.0
0.1
0.2
0.3
0.4
Branch length
(b)
NtPMT1
1
NtPMT2
2
3
4
5
Taxa
6
7
8
9
10
0.0
0.1
0.2
0.3
0.4
Branch length
(c)
NtMPO1
1
2
3
4
Taxa
5
6
7
8
9
0.0
0.05
0.1
0.15
0.2
0.25
Branch length
Supplementary Figure 4. Molecular phylogenetic trees of genes on the pyrrolidine ring pathway
(a) Ornithine decarboxylase (ODC). (b) Putrescine N-methyltransferase (PMT). (c) N-methylptoresine oxidase (MPO).

### Slide 5
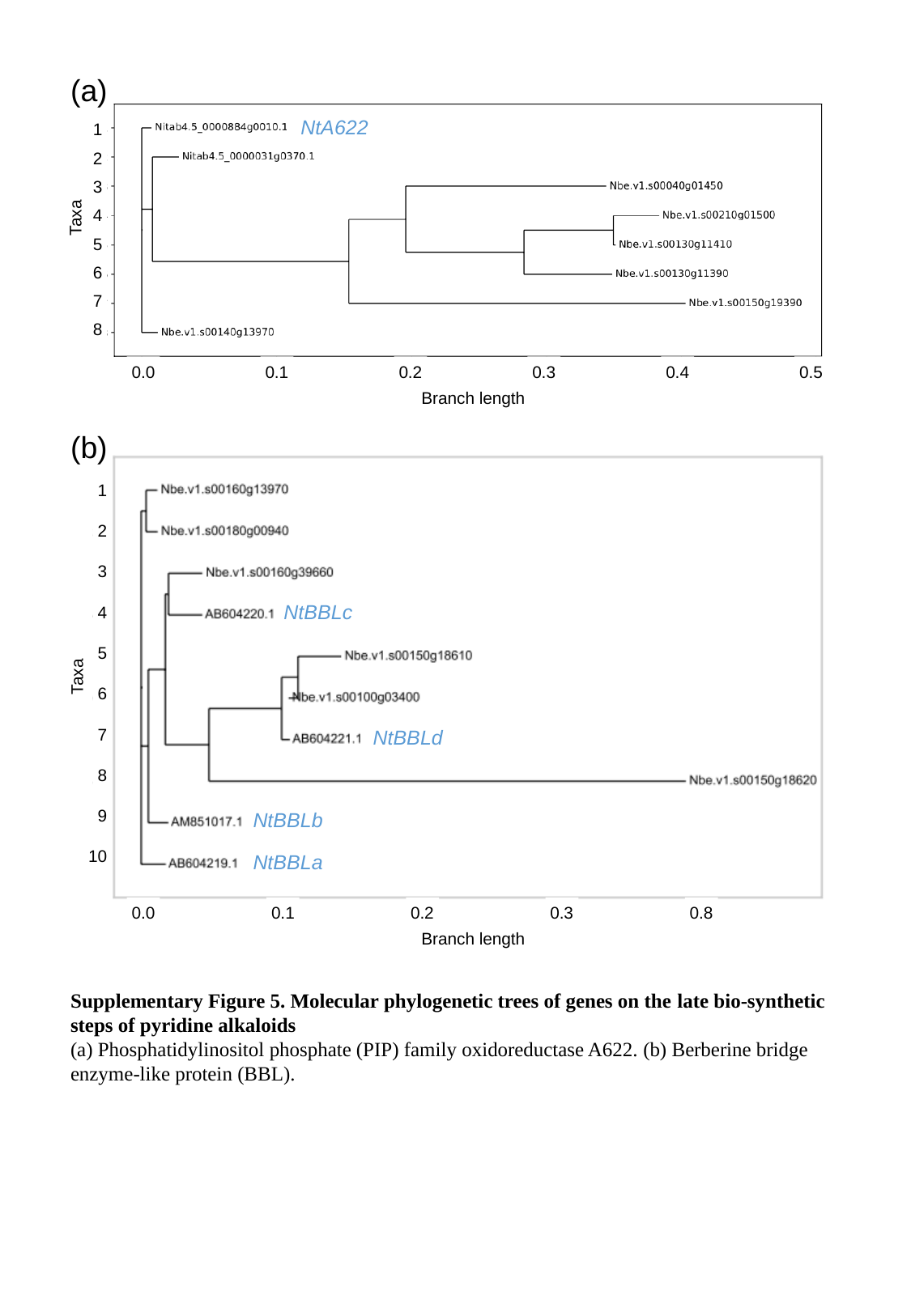

(a)
NtA622
1
2
3
4
Taxa
5
6
7
8
0.0
0.1
0.2
0.3
0.4
0.5
Branch length
(b)
1
2
3
NtBBLc
4
5
Taxa
6
NtBBLd
7
8
9
NtBBLb
10
NtBBLa
0.0
0.1
0.2
0.3
0.8
Branch length
Supplementary Figure 5. Molecular phylogenetic trees of genes on the late bio-synthetic steps of pyridine alkaloids
(a) Phosphatidylinositol phosphate (PIP) family oxidoreductase A622. (b) Berberine bridge enzyme-like protein (BBL).
